# Meteorin resolves nociceptive hypersensitivity by reducing connexin-mediated coupling in satellite glial cells

**DOI:** 10.1101/2025.06.16.659847

**Authors:** Lone Tjener Pallesen, Anna-Kathrine Pedersen, Ishwarya Sankaranarayanan, Rachel Feldman-Goriachnik, Ole Andreas Ahlgreen, Alana Miranda Pinheiro, Jonas Baake, Mads Würgler Hansen, Mette Richner, Laurits Dall Ellegaard, Rachele Rossi, Kenneth A. Petersen, Lone Fjord-Larsen, Jesper V. Olsen, Gordon Munro, Menachem Hanani, Theodore J. Price, Christian Bjerggaard Vægter

## Abstract

Neuropathic pain, a persistent condition arising from injury to the nervous system, involves complex interactions between neurons and non-neuronal cells, including satellite glial cells (SGCs) in the dorsal root ganglia (DRG). In this study, we examined the glial-targeting effects of meteorin, a neurotrophic protein with gliogenic properties, using mouse models of neuropathic and inflammatory pain. Systemic meteorin administration reversed mechanical hypersensitivity across diverse neuropathic and inflammatory pain models, with therapeutic effects persisting beyond the treatment period. We identified SGCs as the principal site of meteorin expression and action in the DRG, where it selectively activated SGCs and altered their functional state. Proteomic profiling revealed meteorin-mediated downregulation of gap junction proteins in SGCs, particularly connexin 43, which was corroborated by immunohistochemical analyses. Functional assessments demonstrated that meteorin treatment normalized injury-induced increases in intercellular coupling between SGCs, establishing a mechanistic link between glial network modulation and pain resolution. These findings identify meteorin as a regulator of SGC communication through connexin-dependent mechanisms. The sustained therapeutic effects and multi-model efficacy highlight meteorin as a potential intervention for neuropathic pain while advancing our understanding of SGC plasticity in sensory processing.

## Introduction

Neuropathic pain, a persistent and often intractable form of pain, results from injuries or diseases affecting the nervous system. Characterized by a complex set of symptoms including allodynia, hyperalgesia, and spontaneous pain, its management and treatment remain a daunting task for healthcare professionals, often leading to significant challenges and burdens for patients^1–3^.

The mechanisms underlying neuropathic pain are multifaceted, involving both neuronal and non-neuronal cells in the peripheral and central nervous systems^4^. Central to advancing our understanding of neuropathic pain is the role of satellite glial cells (SGCs) in the dorsal root ganglia (DRG). These cells form a tight sheath around sensory neuron cell bodies and play a crucial role in maintaining the neuronal microenvironment. Recent advances in the field have highlighted the dynamic nature of SGCs, particularly their responsiveness to neuronal stress and injury. These responses include changes in SGC-SGC and SGC-neuron communication, contributing to the pathophysiology of neuropathic pain and making them potential targets for therapeutic interventions^5^. However, the detailed mechanisms behind these changes in SGC behavior, and their full impact on pain outcomes are not yet fully explored, representing a significant gap in our current knowledge. A better understanding of SGC function and regulation could provide valuable insights into neuropathic pain mechanisms and reveal new therapeutic targets.

Meteorin is a 30 kDa secreted protein expressed in neural progenitors and the glial cell lineage during the development of both the central and peripheral nervous systems. It has been shown to selectively promote astrocyte formation from mouse cerebrocortical neurospheres in differentiation cultures, while also inducing the maturation of cerebellar astrocytes. As neuronal tissues develop, meteorin expression becomes enriched in glial cells^6^. In the mature nervous system, meteorin mRNA is detectable in glial cell types such as Bergmann glia of the cerebellum^7,8^, Müller glia of the retina^9,10^ and SGCs in the PNS^11–16^. Furthermore, SGCs have been identified as targets of meteorin, which induces morphological changes in these cells^6^. Notably, while meteorin does not directly target sensory neurons, conditioned media from meteorin-stimulated SGC cultures can promote neurite outgrowth in cultured DRG sensory neurons^6^ .

Interestingly, studies have demonstrated that meteorin modulates pain outcomes in rodent models^17–20^. Investigating the interaction between meteorin and SGCs thus provides an opportunity to uncover mechanisms underlying neuropathic pain and to explore potential avenues for therapeutic intervention. In this study, we examined the effects of meteorin on SGCs within the DRG in the context of nociceptive sensitization relevant to neuropathic pain. We focused on meteorin’s potential to alter SGC behavior, particularly in relation to their coupling and inter-cellular communication known to be critical for pain modulation. Our findings suggest that meteorin influences SGC-SGC coupling by modulating the expression or function of connexin 43 (Cx43), highlighting SGCs as promising targets for future pain relief strategies.

## Results

### Meteorin Alleviates Nociceptive Hypersensitivity in Mouse Models of Neuropathic and Inflammatory Pain

Previous studies have demonstrated meteorin’s ability to reduce mechanical allodynia in rat chronic constriction injury (CCI) models^17,19^ and mouse paclitaxel-induced peripheral neuropathy models^18^, and hyperalgesic priming^20^. In this study, we expanded our investigation of meteorin’s anti-allodynic effects in additional murine pain models.

We first employed the spared nerve injury (SNI) model. Unlike the CCI model, which involves light ligation of the sciatic nerve leading to intraneural edema and inflammation, the SNI model entails partial sciatic denervation, resulting in significant and prolonged changes in behavioral measures of mechanical sensitivity^21,22^. We observed the development of robust mechanical allodynia within 2 days post-surgery (**Figure 1**). Administration of 5 injections of meteorin from Day 5 to 15 resulted in a significant reversal of mechanical allodynia, persisting several weeks throughout the testing period following the final treatment.

**Figure 1.**
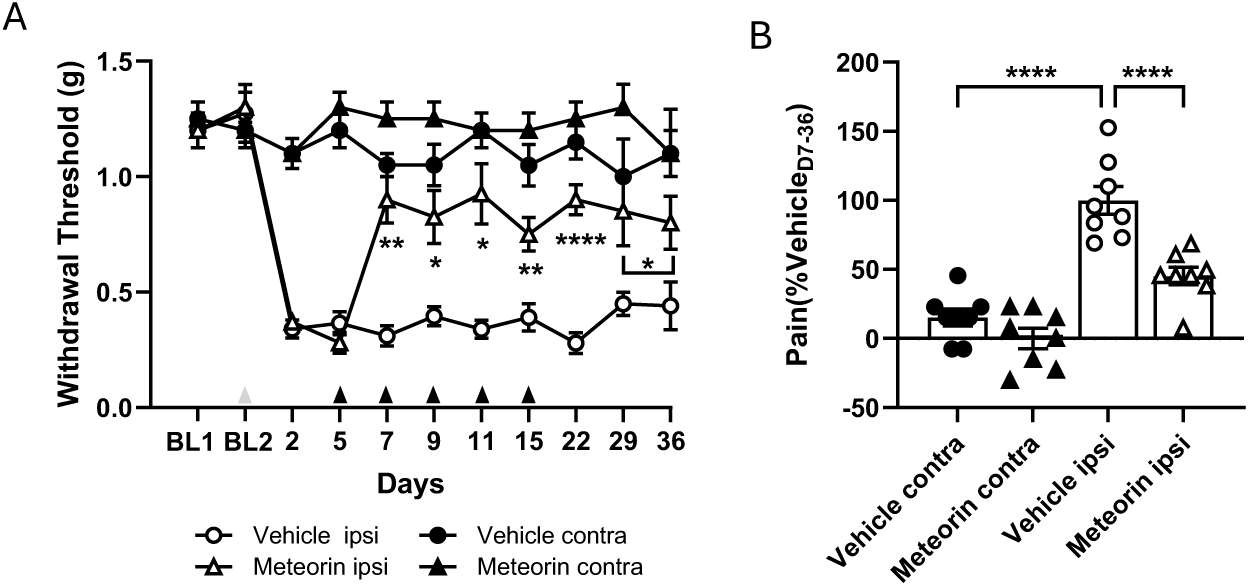
Meteorin treatment alleviates mechanical allodynia in SNI mice. **A)** Hindpaw withdrawal thresholds to ipsilateral (ipsi) and contralateral (contra) stimulation with von Frey filaments were assessed before and after SNI surgery (grey arrowhead) at baseline (BL1 + BL2) and regularly thereafter from Day 2 until Day 36. Meteorin (1.8 mg/kg, s.c.) or vehicle (D-PBS, 20 ml/kg, s.c.) was injected at Days 5, 7, 9, 11, and 15 (black arrowheads) and revealed a significant effect of meteorin treatment, F(3,28)=115.3, p<0.0001 Two-way RM ANOVA. *p<0.05, **p<0.01, ****p<0.0001 versus Vehicle ipsi, Tukey’s test. **B)** Comparison of treatment effects are presented as Pain (% of Vehicle, area under curve for Day 7-36) for each treatment and normalized to the ipsilateral Vehicle treatment. F(3,28)=33.29, p<0.0001, One way ANOVA followed by Tukey’s test; ****p<0.0001 versus Vehicle ipsi. Data are presented as mean ± SEM. All groups n=8, except at Days 29 and 36 for which all groups n=4. No change in the contralateral paw withdrawal threshold was observed.

We also assessed meteorin’s efficacy in mice with neuropathic pain induced by the chemotherapeutic drug cisplatin. Our recent findings indicated that repeated injections of meteorin can fully reverse mechanical hypersensitivity in mice with another chemotherapeutic drug, paclitaxel^18^. Following the initial cisplatin administration, mice developed robust neuropathic hypersensitivity by Day 11, as indicated by a marked reduction in withdrawal thresholds from baseline (**Figure 2**). Even after a second cycle of cisplatin treatment maintaining low mechanical thresholds, subsequent meteorin treatment starting at Day 18 significantly reversed mechanical hypersensitivity from Day 22 to Day 32 (P<0.0001 versus cisplatin + vehicle). From approximately Day 43, mechanical thresholds began to spontaneously resolve in cisplatin-treated mice with vehicle administration.

**Figure 2.**
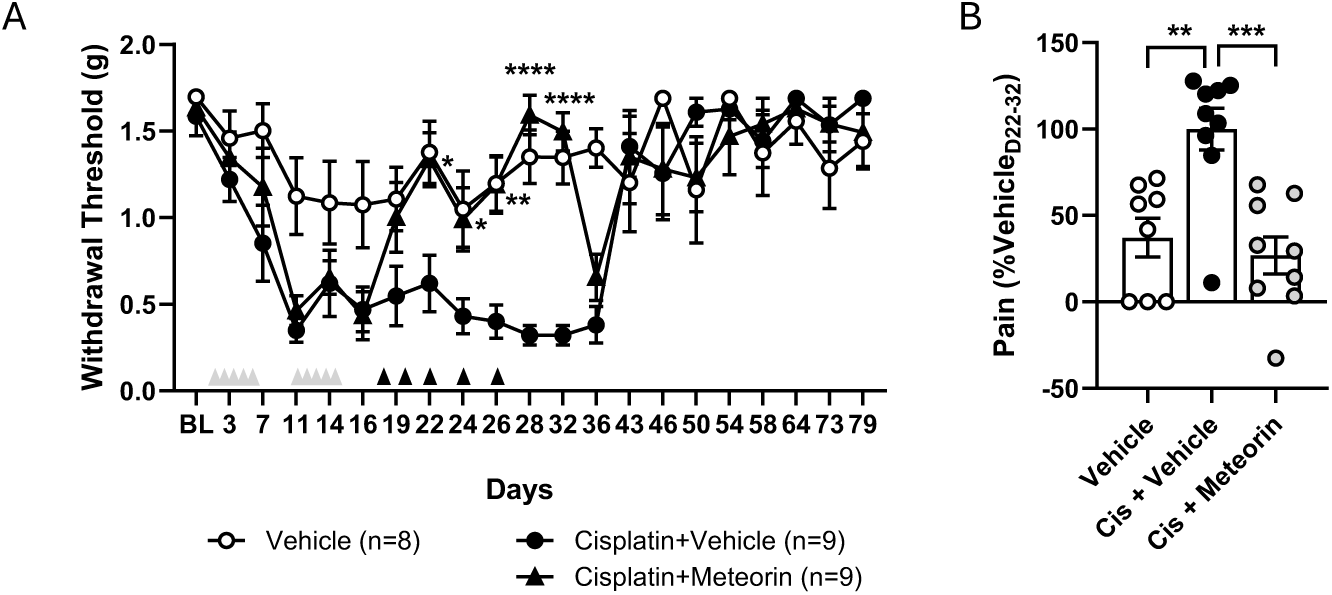
Meteorin treatment reverses mechanical allodynia in cisplatin treated mice. **A)** Hindpaw withdrawal thresholds to stimulation with von Frey filaments were assessed at baseline (BL), during two cycles of cisplatin treatments (grey arrowheads) and regularly thereafter from Day 16 until Day 79. Meteorin (1.8 mg/kg, s.c.) or vehicle (D-PBS, 20 ml/kg, s.c.) was injected at Days 18, 20, 22, 24, and 26 (black arrowheads) and revealed a significant effect of meteorin treatment F(3,31)=10.39, p<0.0001; ANOVA followed by Tukey’s test, *p<0.05, **p<0.01, ****p<0.0001 vs Cisplatin + Vehicle. **B)** Comparison of treatment effects presented as Pain (% of Vehicle, area under curve for Day 22-32) for each treatment and normalized relative to the Cisplatin + Vehicle treatment. F(3,31)=8.940, p=0.0002, One way ANOVA followed by Tukey’s; **p<0.01***, p<0.001 vs Cis + Vehicle. Data are presented as mean ± SEM. All groups n=9, except Vehicle n=8.

Based on these results (**Figures 1 and 2**), combined with previously published data^17–19^, we conclude that meteorin treatment effectively reverses neuropathic hypersensitivity across several rodent models of peripheral nerve injury and chemotherapy-induced neuropathy. However, the efficacy of meteorin in inflammatory pain models had not been previously investigated. Therefore, we employed the Complete Freund’s Adjuvant (CFA) model of inflammatory hyperalgesia. Five days post-CFA injection, all mice exhibited reduced mechanical withdrawal thresholds which was significantly reversed by systemic administration of meteorin (**Figure 3A-B**). At the end of the study, CFA mice were administered the partial µ-opioid receptor agonist buprenorphine to establish assay sensitivity to a known analgesic drug. As expected, buprenorphine produced a marked increase in withdrawal thresholds (P<0.0001 vs vehicle) (**Supplementary figure S1**). An almost 2-fold increase in hindpaw width 5 days post-CFA injection confirmed a robust inflammatory response in CFA mice (**Figure 3C**), consistent with the observed low mechanical withdrawal thresholds. Notably, hindpaw widths remained unaffected by meteorin treatment, suggesting that the reversal of CFA-mediated hyperalgesia did not occur through an overt anti-inflammatory mechanism.

**Figure 3.**
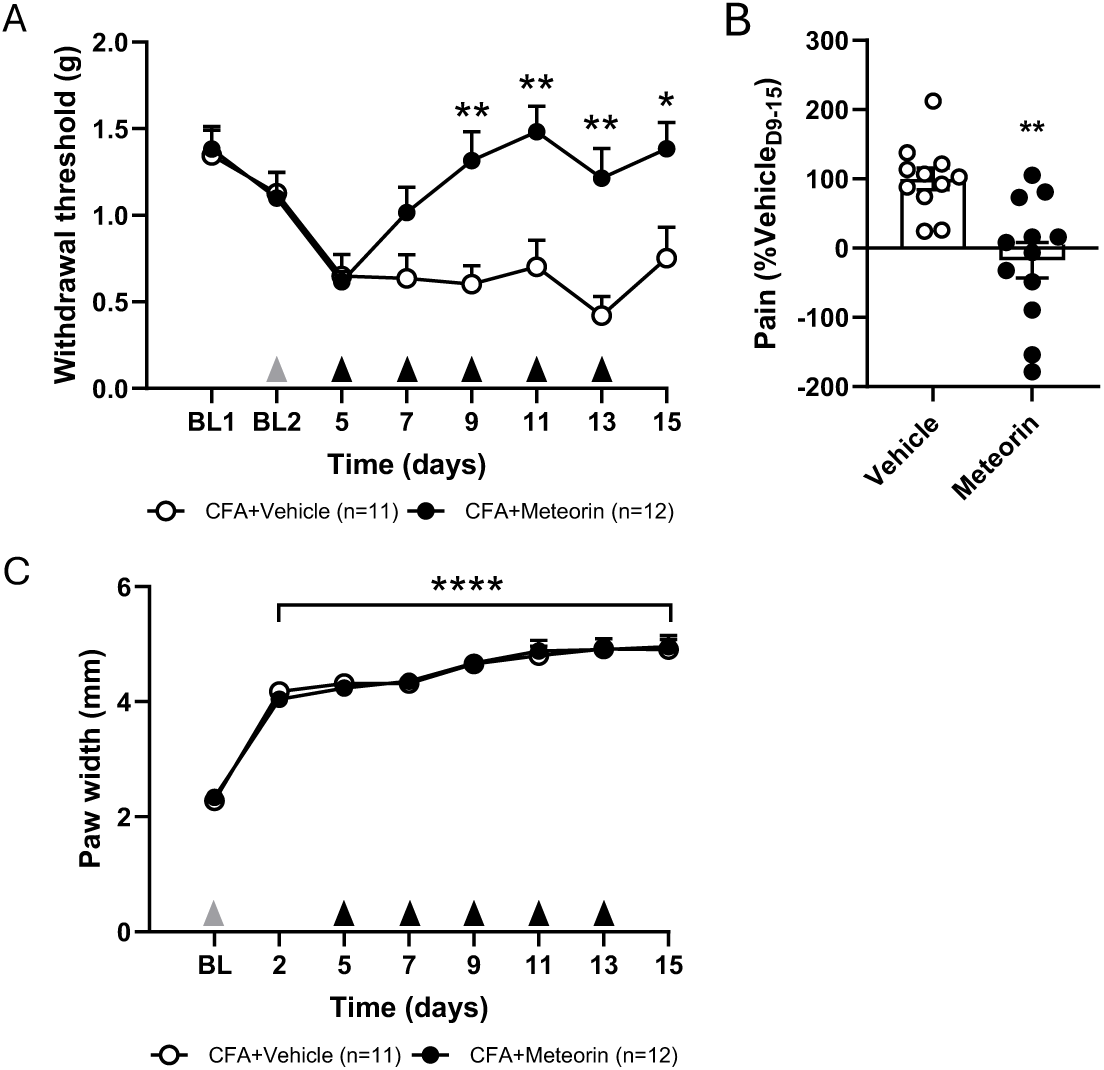
Meteorin treatment reverses inflammatory hyperalgesia in CFA mice. **A)** Hindpaw withdrawal thresholds to stimulation with von Frey filaments were assessed at baseline (BL) and then 5 days after hindpaw injection of complete Freund’s adjuvant (grey arrowhead) to confirm the presence of inflammatory hyperalgesia. Meteorin (1.8 mg/kg, s.c.) or vehicle (D-PBS, 20 ml/kg, s.c.) was injected at Days 5, 7, 9, 11, and 13 (black arrowheads) revealing a clear effect of meteorin treatment F(1,21)=25.31, p<0.0001; Two-way RM ANOVA followed by Tukey’s test, *p<0.05, **p<0.01, vs Vehicle. **B)** Comparison of treatment effects presented as Pain (% of Vehicle, area under curve for Day 9-15) and normalized to vehicle treatment, Mann-Whitney test, **p<0.01 vs Vehicle. **C)** Paw width (mm) measured as a surrogate marker of inflammatory edema revealed a significantly increased paw width in both vehicle and meteorin treated CFA mice, F(7,88)=51.39, p<0.0001, Two-way RM ANOVA followed by Tukey’s test ****p<0.0001 versus baseline. Data are presented as mean ± SEM. Vehicle group n=11, meteorin group n=12.

### Meteorin Targets Satellite Glial Cells

Previous studies have indicated that meteorin can stimulate astrocytic gliogenesis^6,23,24^ and may act as a negative regulator of reactive gliosis^25^. Given that meteorin is a 30 kDa secreted protein, unlikely to cross the blood-brain barrier (BBB) in significant amounts^17^ , we hypothesized that exogenously administered meteorin targets peripheral cells. A prior study demonstrated that meteorin activated cultured SGCs, while DRG neurons remained unaffected. Furthermore, conditioned media from meteorin-treated SGCs (but not from untreated controls) induced neurite outgrowth, suggesting a direct action of meteorin on SGCs, with secondary neuronal effects mediated by activated SGCs^6^. This supports the idea that endogenous meteorin plays a crucial role in neuronal processes and SGC-neuron communication within the DRG. Our single cell RNAseq of lumbar mouse DRGs^14^ showed meteorin expression mainly in SGCs, and to a minor degree in Schwann cells (**Figure 4A**). This expression pattern is in accordance with our previous transcriptional analysis of FACS-sorted SGCs^15^ as well as other available single nucleus/cell transcriptional analyses^11–13,16^. To further validate meteorin expression in SGCs, we performed RNAscope to visualize mRNA transcripts on DRG sections in combination with immunohistochemistry (IHC) of the SGC marker FABP7. RNAscope confirmed meteorin expression mainly in SGCs (**Figure 4B**). We attempted to visualize meteorin expression in SGCs by IHC. Although we detected meteorin in cerebellar Bergmann glia (**Supplementary figure S2**), in accordance with Jørgensen *et al.*^7^, we were unable to visualize meteorin protein in the DRG.

**Figure 4.**
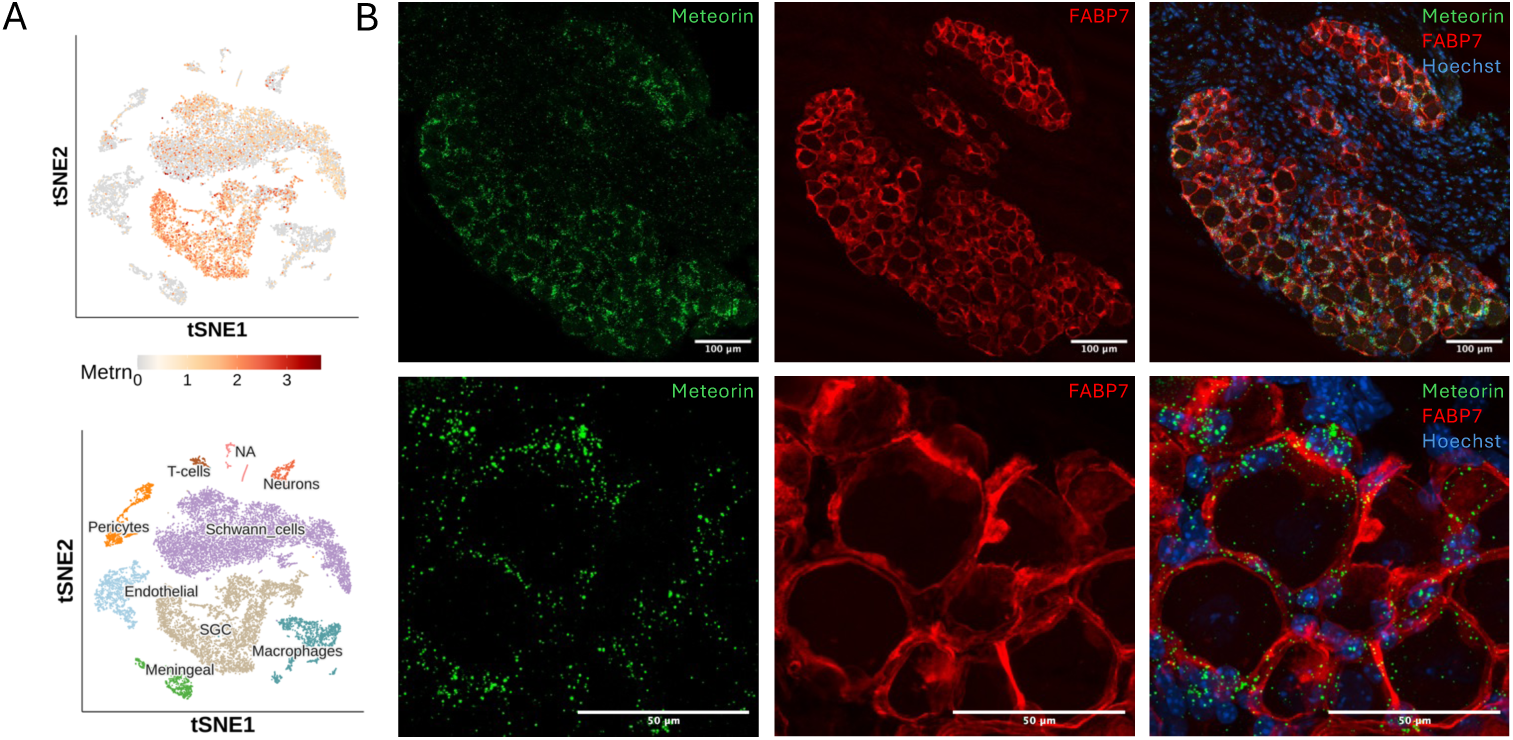
Meteorin expression in the mouse dorsal root ganglion (DRG) **A)** Clustering and annotation of cell populations in the DRG. Classifications of cell populations in lumbar mouse DRGs by single cell RNAseq. T-SNE plot visualizing the cellular heterogeneity of DRG cells with 9 distinct cell clusters. Dots, individual cells; colors, individual cell types as labelled. Top t-SNE plot demonstrate meteorin expression in the DRG mainly by SGCs. **B)** RNAscope of mouse DRG visualizing meteorin mRNA transcripts in SGCs. Integrated co-detection workflow for meteorin mRNA and IHC for FABP7 to visualize SGCs. Confocal microscopy of a lumbar DRG stained with probes targeted to meteorin mRNA (green), antibody against FABP7 (red) and DAPI (blue). Top: 10x magnification. Bottom: 40x magnification.

SGCs, and not neurons, have previously been proposed as meteorin targets in the DRG^6^, suggesting an auto-/paracrine effect of endogenous meteorin on SGCs. To verify SGCs as meteorin target cells, we stimulated SGC-enriched mouse DRG cultures with meteorin and assessed potential changes in the activation of common cellular downstream signaling proteins Akt and ERK, as described previously by others^6,23,25^ . Stimulation with meteorin 800 ng/ml for 15 minutes induced a significant phosphorylation of Akt (Thr308) (42% +/-9% relative to unstimulated control, p=0.007) (**Figure 5A-B**). We also observed a slight increase in phosphorylation of ERK2 (19% +/-1.4%, p = 0.0003) relative to unstimulated control, whereas there was no change in ERK1 (8% +/- 11%, p = 0.23).

**Figure 5.**
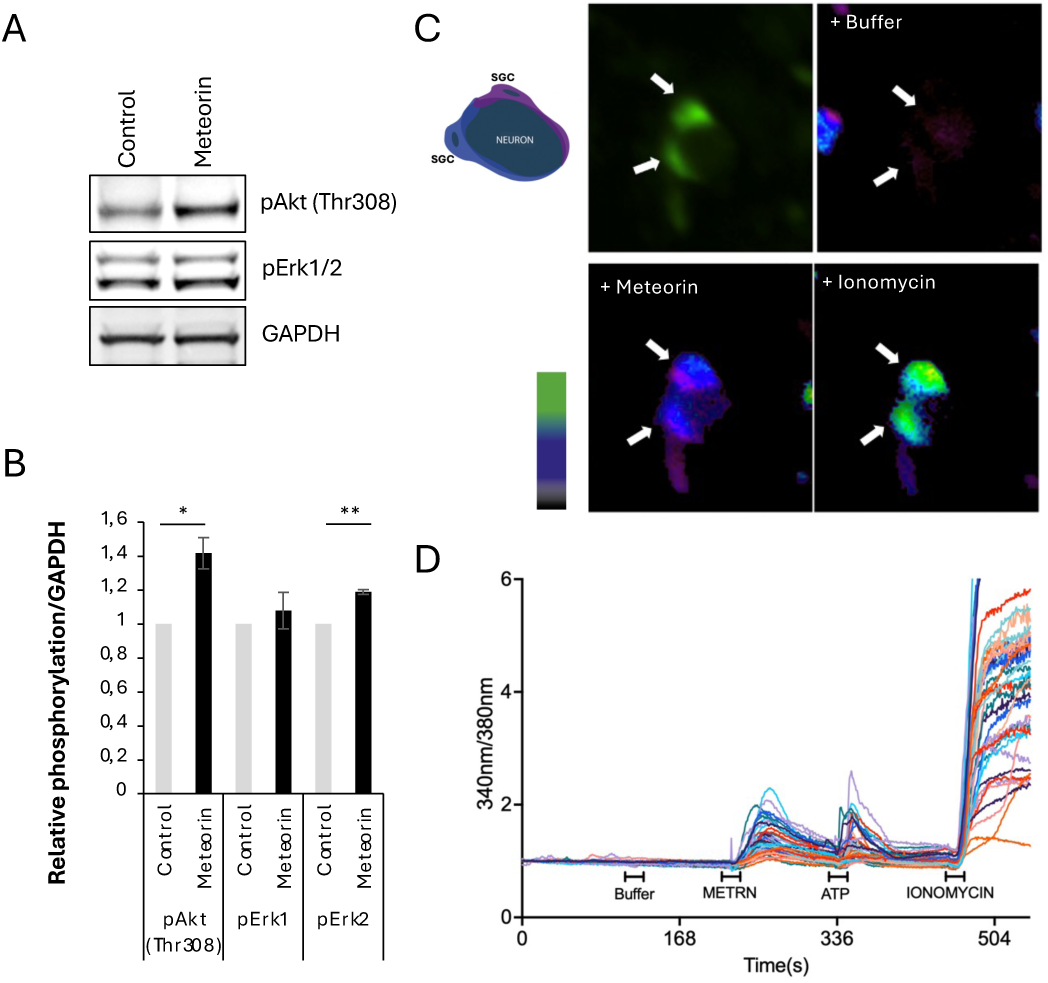
Meteorin stimulation of primary cultures of mouse DRGs. **A)** Representative western blots showing SGC-enriched cultures incubated with or without 200 ng/ml meteorin for 15 min. The level of phosphorylation of ERK1/2(Thr202/Tyr204) and Akt(Thr308) as well as protein levels of GAPDH were detected by western blotting. **B)** pAkt and pERK1/2 levels in stimulated SGC cultures quantified by Western blot and normalized to control. One-way RM ANOVA, n = 4, *p<0.05, **p<0.01. **C)** Calcium response assay. Loading mouse DRG cultures with the calcium sensitive indicator Fura2-AM allows evaluation of responsiveness to stimulation by meteorin, ATP and ionomycin. Fura2-AM preferentially stains SGCs, here indicated by white arrows. Insert: Illustration of the SGC-neuron unit illustrated by Fura2-AM assay. **D)** The fluorescence ratio 340nm/380nm indicate SGC and neuron Ca^2+^ signaling after the addition of the indicated stimulant; meteorin, ATP or ionomycin.

Although SGC-enriched primary cultures can be prepared in reasonable quantity and purity to allow SGC-related investigations^26–30^, this system has limitations. Primarily, SGCs *in vivo* exist in close proximity to the neuronal soma in distinct SGC-neuron units, defined by several SGCs intimately enclosing individual (or occasionally a few) neuronal soma^5^. Furthermore, it has been reported that SGCs may undergo rapid dedifferentiation upon losing contact with the neuron^31^, eventually regressing into a precursor state converging with that of dedifferentiated cultured Schwann cells^32^. To verify SGCs as meteorin targets under conditions closer to their fully differentiated *in vivo* state, we used short-term cultured (18 hours) SGC-neuron units from adult mouse DRGs, with SGCs identified based on their cellular characteristics closely enveloping the neuronal soma (**Figure 5C**). The cultures were loaded with the calcium indicator Fura2-AM, which primarily loaded SGCs and to a lesser degree the neurons (**Figure 5C**). Upon stimulation with meteorin, we observed a calcium elevation in a majority of the SGCs (**Figure 5C-D**). A total of 719 SGCs in 7 independent replicates were analyzed, of which 53% responded to meteorin stimulation. In contrast, >95% of SGCs responded to ATP, which was included as a positive control since the SGC calcium response to ATP has been well described by others^33–36^. In our analysis, we only included SGCs that were responsive to ionomycin as a final stimulant, verifying cell viability (**Figure 5D**). These data support SGC reactivity to meteorin but may also suggest SGC heterogeneity regarding the yet unidentified meteorin receptor.

These findings, along with the previously published data by Nishino *et al.*^6^, indicate that SGCs are the primary targets of meteorin, but it can be argued that the SGC calcium response could be a secondary effect of neuronal stimulation and signaling. Neuronal depolarization with KCl has been shown to promote both a neuronal response and an SGC response, the latter considered secondary to neuron stimulation since SGCs without neuronal contact were unresponsive^37^. To study potential direct meteorin signaling to neurons and the possible secondary induction of an SGC response, we examined a total of 220 individual neuron-SGC units (with a total of 854 SGCs). Only neurons responsive to KCl and only neurons/SGCs responsive to a final ionomycin stimulation were included in the analysis. Approximately 50% of SGCs and 40% of neurons responded to meteorin stimulation; however, neurons responded only when they were surrounded by responsive SGCs, whereas responding SGCs were also observed associated with unresponsive neurons. Chi-square analysis (p=0.013) confirmed a significant correlation between SGC and neuronal responses to meteorin, suggesting that the neuronal calcium response is indeed secondary to SGC activation by meteorin, as reported by others^6^. To further strengthen this conclusion, we investigated the calcium response to stimulation with endothelin 1 (ET1) in a separate experiment with a total of 92 neurons surrounded by 254 SGCs. The cellular response to ET1 has been associated with endothelin receptor A (ETA) for neurons, while endothelin receptor B (ETB) mediates ET1 signaling in SGCs^38–40^. As expected, a large fraction of both SGCs (56%) and neurons (53%) were responsive to endothelin in our analysis. However, in contrast to the results obtained with meteorin, we did not observe any correlation between the signaling of neurons and associated SGCs (p=0.66), confirming that SGCs and neurons responded independently to ET1.

### Meteorin stimulation induces changes in the DRG proteome

To elucidate the cellular response underlying meteorin’s pain-resolving effect, we investigated meteorin-induced changes in protein expression levels using mass spectrometry (MS)-based proteome analysis of SGC-enriched DRG short-term cultures. The cultures were incubated for 48 hours with or without 200 ng/ml meteorin, followed by cell lysis and protein digestion for quantitative analysis by liquid chromatography-tandem mass spectrometry (LC-MS/MS). The proteome analysis identified more than 8,000 proteins. Principal component analysis and unbiased hierarchical clustering of all identified proteins revealed that meteorin-stimulated and control samples clustered according to treatment after data normalization (**Supplementary figure S3**). Correspondingly, we observed a significant response to meteorin in terms of altered protein expression patterns, as visualized in the volcano plot in **Figure 6A**. These data indicate that the DRG is a relevant meteorin target in the peripheral nervous system (PNS). Among the upregulated proteins, we found several associated with neuronal cell growth, differentiation, and neurite outgrowth (**Supplementary Table 1**), aligning with previous findings^6^.

**Figure 6.**
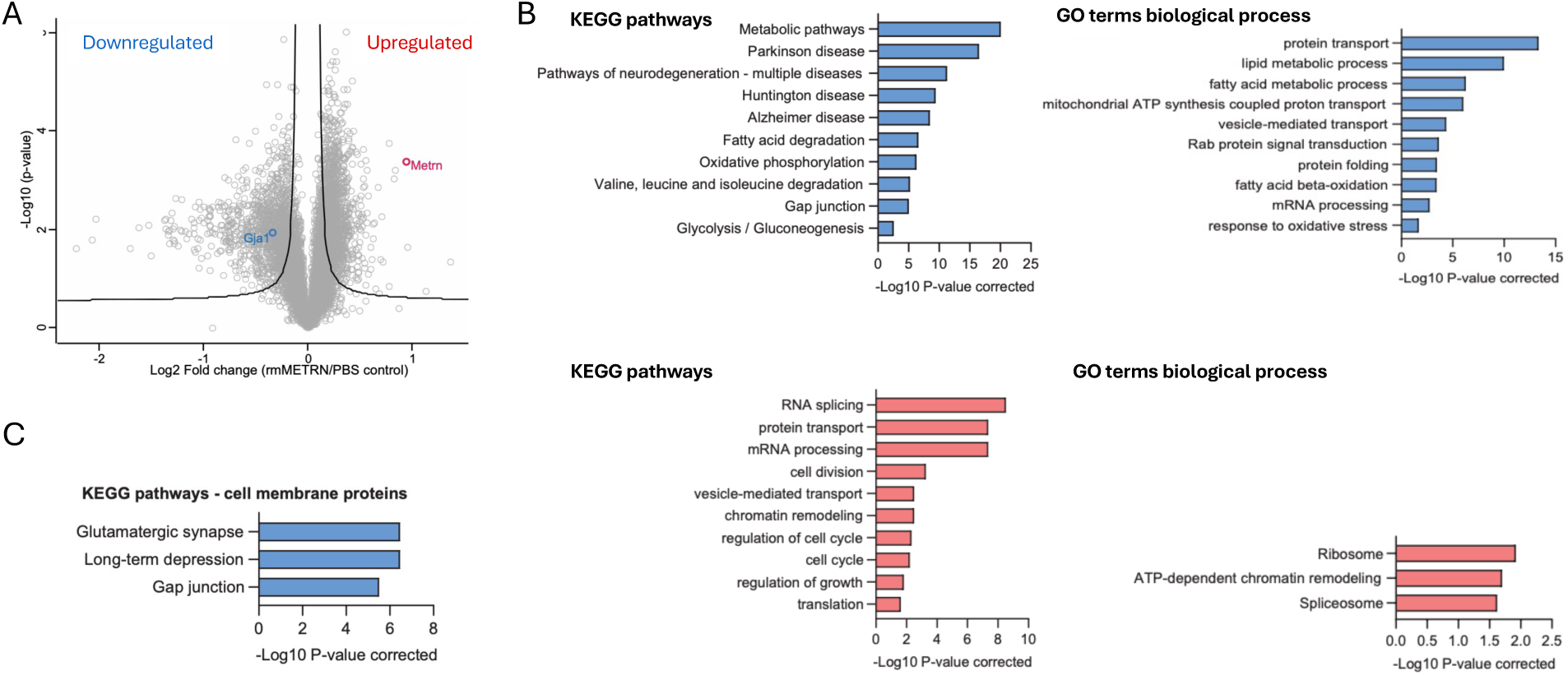
Global proteome changes of meteorin stimulated DRG cultures. **A)** Volcano plot illustrating proteins regulated by meteorin. Volcano plot generated based on t-test and illustrating proteins significantly regulated by meteorin as compared to PBS determined by a hyperbolic curve threshold (FDR<0.05 and S0=0.1). The plot shows the –log(p-value) over the log2 fold change in protein levels between meteorin and PBS. **B)** Enriched GO terms and pathways for meteorin-regulated proteins. The figures show selected among the top significantly (corrected p-value < 0.05) enriched GO (gene ontology) terms for biological process and KEGG (Kyoto Encyclopedia of Genes and Genomes) pathways, associated with up- and down-regulated proteins, respectively. Terms/pathways in red color refer to upregulated proteins and blue color refer to downregulated proteins. **C)** KEGG pathway enrichment analysis for meteorin-downregulated membrane proteins identified using DeepLoc-predicted subcellular localization.

Meteorin itself was among the proteins showing the largest fold change when comparing meteorin-stimulated and control samples, validating its presence in the stimulated cultures after 48 hours (**Figure 6A and Supplementary Table 1**). Gene ontology (GO) and pathway enrichment analysis revealed various significantly enriched KEGG pathways and GO terms associated with meteorin-regulated proteins. For meteorin-downregulated proteins, significantly overrepresented KEGG pathways and GO biological processes included metabolic and fatty acid processes, neurodegenerative diseases, and gap junctions. For upregulated proteins, significantly enriched GO biological processes and KEGG pathways included processes related to protein transport, RNA, and cell cycle (**Figure 6B**). Given that proteins localized in the cell membrane are central in mediating the actions of secreted proteins like meteorin, as well as pain responses in general, we specifically focused on the identified regulated membrane proteins using the DeepLoc-predicted subcellular localization (see methods). KEGG pathway enrichment analysis for meteorin-downregulated membrane proteins demonstrated prominent overrepresentation of gap junctions, emphasizing their potential importance in mediating the effects of meteorin (**Figure 6C**). Furthermore, for unbiased identification of regulated proteins implicated in pain, we utilized the “Diseases” database of disease–gene associations mined from literature and data integration, focusing on terms related to pain (including “pain disorder”, “complex regional pain syndrome”, and “myofascial pain syndrome”)^41^. This analysis resulted in a list of 18 proteins, which are displayed in a STRING-based network in **Supplementary Figure S3D**.

### Meteorin Reverses Injury-Induced Upregulation of connexin 43 and SGC Coupling

Intriguingly, KEGG pathway enrichment analysis revealed gap junctions as one of the significantly enriched cellular pathways associated with meteorin-downregulated proteins (**Figure 6B-C**). Among the downregulated proteins related to gap junctions, we identified the gap junction alpha-1 protein, connexin 43 (Cx43) (**Figure 6A**). It is well established that gap junctions facilitate long-range interactions in sensory ganglia^5,42^. After nerve injury, gap junctions are upregulated, and injury-associated pain hypersensitivity behavior can be reduced by the gap junction blocker carbenoxolone, suggesting an important role for SGC-SGC coupling via gap junctions in conditions of neuropathic pain (reviewed in ^5,43^). Our data confirm an increased expression of Cx43 in the DRG of cisplatin-treated mice. This increase was mitigated by meteorin administration compared with vehicle in cisplatin-treated mice (**Figure 7A**).

**Figure 7.**
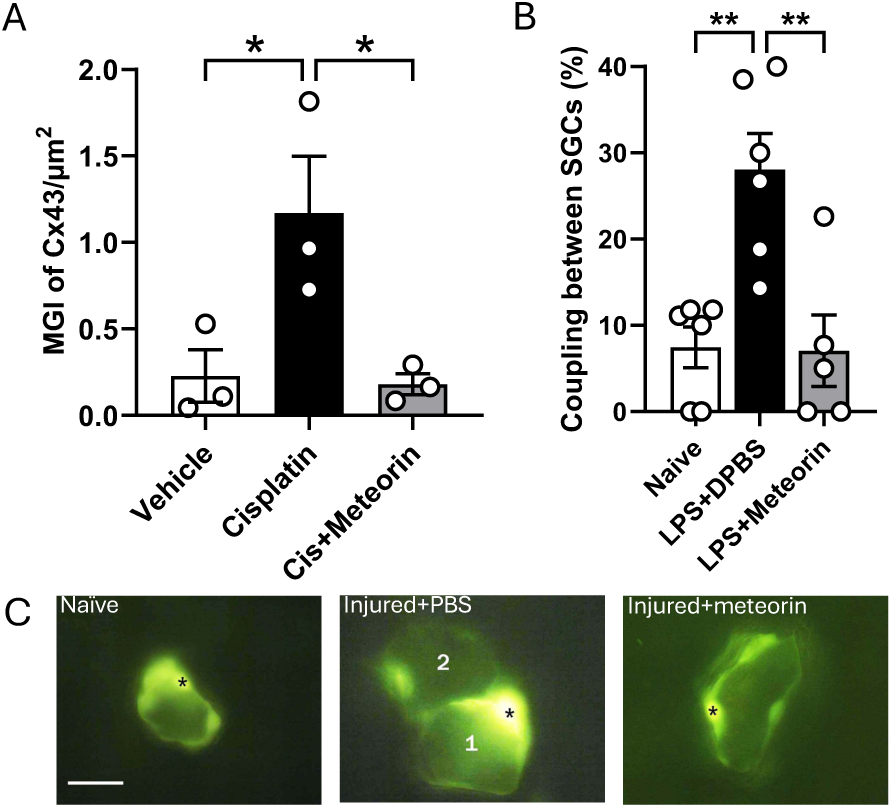
Meteorin reverses Connexin 43 (Cx43) upregulation and SGC coupling induced by injury. **A)** DRGs were obtained at Day 34 from cisplatin-treated mice and immunoreactivity to Cx43 assessed and presented Mean Grey Intensity (MGI) of DRG Cx43 expression per DRG μm^2^. Cisplatin-induced upregulation of Cx43 was reversed by meteorin treatment, F(2,6)=6.868, p=0.0281, One Way ANOVA followed by Tukey’s test, *P<0.05 vs Vehicle. **B)** SGC coupling mediated by LPS injection in mice is reduced by meteorin treatment **P<0.01 Mann-Whitney test. Data are shown as means ± S.E.M. <u>Intracellular</u> Luficer yellow labeling and recording in SGCs in mouse DRG, injected with Lucifer yellow dye. Asterisks indicate the injected SGCs. **C)** Pictures from left to right show a naïve mouse, an injured mouse (numbers indicate two separate neurons) treated with PBS, and an injured mouse treated with meteorin. Calibration is 20 μm.

To further study the functional consequences of meteorin-induced reversal of Cx43 expression, we next studied the extent of SGC-SGC coupling. We have previously demonstrated that the fluorescent dye Lucifer yellow, when injected into one SGC, spreads to other SGCs surrounding the same neuronal soma. Furthermore, in a wide range of models involving sensitization of nociceptive pathways, increased coupling among the SGCs not only intensified within the sheaths surrounding individual neurons but also extended to sheaths surrounding adjacent neurons^42,44–52^. Recently, we have shown that meteorin treatment blocks the hyperalgesic priming actions of the cytokine IL-6 in mice^20^. Accordingly, we administered mice intraperitoneally with LPS which causes a systemic release of pro-sensitizing cytokines to induce sustained increase in SGC dye coupling^47,48^. LPS administered mice were injected with meteorin or PBS every second day from Day 2 to Day 14. Lumbar DRGs were extracted on Day 18, and individual SGCs were subsequently injected with Lucifer yellow using sharp glass microelectrodes. We found that meteorin reduced SGC coupling in LPS sensitized mice 4-fold (P<0.01 vs LPS alone) as shown in **Figure 7B**. A similar trend of meteorin treatment was noted in SNI mice (**Figure 7C** and **Supplementary Figure S4**), albeit the limited number of injured neurons per DRG in the SNI model relative to the systemic LPS-model precluded more robust interrogation. Collectively, our data demonstrate that meteorin treatment reversed the increased Cx43 expression and SGC-SGC coupling induced by inflammation or cisplatin, likely underlying its observed analgesic effects.

## Discussion

In this study, we investigated the role of meteorin in modulating SGC activity and its implications for neuropathic pain. We found that meteorin exerts a pronounced effect on SGCs in the DRG, where it downregulates the gap junction protein Cx43. Functional assays further showed that meteorin normalizes the increased coupling between SGCs observed after nerve injury. Systemic administration of meteorin reduced mechanical hypersensitivity in mouse models of both neuropathic and inflammatory pain, with effects persisting beyond the treatment period. Collectively, our results indicate that meteorin contributes to pain relief by acting as a regulator of SGC communication via connexin-dependent mechanisms.

Meteorin is expressed by SGCs within the DRG, and we confirmed that these cells also respond to meteorin. Furthermore, we showed that DRG neurons do not exhibit a calcium response to meteorin unless associated with meteorin-responsive SGCs. This suggests an auto/paracrine signaling mechanism within the PNS, where endogenous meteorin acts rapidly, locally on neighboring SGCs rather than on neurons or distant targets. Paracrine signaling is a common feature in the nervous system, as seen with astrocytes in the CNS, which can release numerous types of signaling molecules that affect nearby neurons, other glial cells, or the astrocytes themselves in auto- and paracrine manners^53–56^. While astrocytes and SGCs are exclusively present in the CNS and PNS, respectively, and structurally appear to be very different, they possess several similar features in their roles of modulating neuronal responses^57,58^. Like astrocytes, SGCs have been reported to release ATP, cytokines, and other signaling molecules that can influence the activity of neighboring SGCs and neurons, contributing to the local cellular environment and potentially modulating pain signaling^5^.

The response of SGCs to neuronal injury or disease has been studied in several rodent models, including nerve injury^15,46,49^, diabetes^59–61^, chemotherapy^18,45^, alpha-herpes viruses^62^, and induced systemic infection (sepsis) by LPS^47,48^. Such studies have convincingly demonstrated activation of SGCs in sensory ganglia with altered expression levels of various proteins such as glial fibrillary acidic protein (GFAP)^45,63–65^, KIR4.1 potassium channels^65–68^, and pro-inflammatory cytokines^69–71^. Increased connexin-mediated SGC-SGC coupling^46,48–50,65,72^, and increased sensitivity to the pain mediator ATP^34,47,73–76^ also appear to be involved in injury-induced SGC activation (reviewed in Hanani and Spray^5^). Yet, most studies are observations of cumulative changes in protein expression, whereas little is known about the acute coupling between neuronal and SGC activity. A pioneering study demonstrated bidirectional communication between SGCs and neurons in short-term cultures, showing how electrical or mechanical stimulation induced calcium signaling in the stimulated cell, which spread to neighboring neurons or glia in a purinergic receptor P2-dependent manner. Furthermore, gap junctions were involved to some extent in the SGC response^77^. Another *ex vivo* study investigated calcium signaling in SGCs and DRG neurons following electrical stimulation. It was found that neuronal stimulation prompted a robust neuronal ATP release, increasing with the frequency of nerve stimulation, and that the released ATP elicited a P2X7-dependent calcium response in the surrounding SGCs. Conversely, the activated SGCs released TNFα, which resulted in increased neuronal excitability^36^. These studies provide evidence for acute bidirectional communication between neurons and their surrounding SGCs. Interestingly, a later *in vivo* study described how a large increase in injury-induced “cross-talk” between adjacent DRG neurons might be explained by neuron-to-SGC-to-neuron communication via Cx43^42^, further supporting bidirectional neuron-SGC communication upon neuron signaling. Indications of independent SGC and neuron signaling also exist. A recent study found evidence of spontaneous SGC activity; investigating *in vivo* SGC calcium activity in transgenic mice using FABP7 to drive expression of the calcium sensor GCaMP6 in SGCs, the authors found that SGCs display spontaneous calcium activity at an arbitrary frequency and that the number of active SGCs increased under conditions of inflammation or nerve injury^78^ (see also Hanstein et al.^75^). Surprisingly, they further found that the spontaneous SGC activity was independent of electrical stimulation of the neurons ^78^, contradicting previous findings by Zhang *et al.*^36^. In the present study, we observed that meteorin can increase the intracellular calcium concentration in SGCs independent of neuronal calcium levels, whereas neuronal activity upon meteorin treatment was only observed when the surrounding SGCs also responded. This result supports the observations by Nishino *et al.*^6^, who found that the neuronal effect of meteorin was dependent on SGC stimulation and likely the release of soluble factors acting on the neurons. Considering the observed analgesic effect of meteorin in several pain models, our results suggest that meteorin-stimulated SGCs *in vivo* communicate with nociceptive neurons, although the mechanism of such communication remains to be clarified.

The identity of the meteorin receptor remains elusive, presenting a significant gap in understanding its signaling mechanisms. While HTR2B has been proposed as a candidate^79,80^, single-cell transcriptional analyses suggest that it is not expressed in relevant DRG cell types^11–16^. Recent studies have used *in silico* modeling to predict potential meteorin receptors. One study indicated that meteorin and meteorin-like are composed of CUB and NTR domains, connected by a hinge region^81^. Though KIT/CD117 was described as a receptor for meteorin-like^82^, structural differences suggest that meteorin may not activate KIT signaling. The identity of the receptor for meteorin remains an important area for further research.

Meteorin’s analgesic effects are noteworthy for their persistence after only a few treatments, suggesting that meteorin has disease-modifying properties. The long-lasting effect indicates that meteorin induces changes at the cellular or molecular level that persist beyond the immediate treatment period. Our study confirms that the expression level and functional SGC-SGC coupling via gap junctions are increased in response to injury, and we demonstrate a reduced coupling by meteorin administration. This relation between the analgesic action of meteorin and its ability to block gap junctions aligns well with previous findings that gap junction blockade or neutralization reduce neuronal excitability and pain behavior^44,45,47,83^ . Transcriptional databases align with the observation that SGCs in mouse DRG mainly express *Gja1* (Cx43), and to a lesser degree *Gjc3* (Cx29)^11–16^. Cx36 has also been described in mouse DRG^84^ but expression in SGCs has not been confirmed by single cell transcriptional analyses^12,85^ . Interestingly, there are no indications that injury or disease increases Cx43 expression at a transcriptional level^11,14,15^. Thus, prior observations on increased expression of gap junctions^46,49,50^, and of Cx43 level^83^ as well as our observations here, might be occurring only at the protein level. In general, connexins have emerged as viable drug targets for a wide range of conditions^86,87^. Although no connexin-specific drugs have received approval yet, the importance of connexins in human physiology and disease suggests that connexin therapeutics are forthcoming. By influencing SGC-SGC coupling, meteorin may be altering the glial environment in a way that reduces pain signaling and contributes to its long-term analgesic effects.

## Methods

### Animals

C57Bl/6JRj mice were purchased from Janvier Labs (France) and ICR (CD-1) mice were purchased from Envigo. Mice were group housed with water and chow available ad libitum in a 12:12 hour light-dark cycle. Spared nerve injury experiments were approved by either 1) the Danish Animal Experiments Inspectorate under the Ministry of Justice (permission no. 2017-15-0201-01192) and carried out according to the European Council directive, and institutional and national guidelines, or 2) the Animal Care and Use Committee of the Hebrew University Hadassah Medical School and conformed to the National Institutes of Health standards for the care and use of laboratory animals. Cisplatin experiments were approved by the Institution Animal Care and Use Committee (IACUC) at the University of Texas at Dallas. Complete Freund’s adjuvant (CFA) experiments were approved by the Danish Animal Experiments Inspectorate, The Danish Veterinary and Food Administration, Ministry of Environment and Food (license no. 2020-15-02001-00438 C1).

### Rodent procedures

Spared nerve injury (SNI): Male and female C57Bl/6JRj mice, 8-14 weeks old, were subjected to the SNI model, and mechanical allodynia was assessed by von Frey testing, as described previously^21,88^. Briefly, the common peroneal and tibial branches of the sciatic nerve were ligated and cut distally to the ligation, just distal to the branching of the sural nerve, which was left untouched. Isoflurane gas (IsoFlo vet, Abbott) was applied with a Univentor 1200 anesthesia unit (Univentor). Analgesia, Lidocaine SAD (10 mg/ml; Amgros I/S) applied on wound; temgesic, buprenorphine (0.3 mg/ml; RB Pharmaceuticals); and antibiotics, pentrexyl (250 mg/ml; Bristol-Myers Squibb). Temgesic and pentrexyl were mixed and diluted 1:10 in isotonic saline (9 mg/ml; Fresenius Kabi) and were injected subcutaneously (s.c.) at 0.1 ml. Recombinant mouse meteorin (AF3475, R&D systems) was injected s.c. over the shoulders in the skin at 1.8 mg/kg in 0.5 ml vehicle (saline) at Days 5, 7, 9, 11 and 15 after SNI surgery.

CFA inflammatory hyperalgesia: Complete Freund’s adjuvant (20 ml) was injected s.c. into the dorsal surface of the hind paw of female mice C57Bl/6JRj (Janvier Labs, France) under isoflurane anesthesia. All mice recovered rapidly and were typically active within 5-10 mins upon removal from anesthesia. No post-operative analgesia was provided to help facilitate full development of CFA-induced sensitization. Once hind paw mechanical hypersensitivity was established by Day 5, mice were distributed into 2 separate treatment groups and administered either recombinant mouse meteorin (AF3475, R&D systems) at 1.8 mg/kg (s.c.) or vehicle at Days 5, 7, 9, 11 and 13. Withdrawal thresholds were measured prior to drug injection on each day. At the end of the study, 6 female CFA administered mice repeatedly treated with vehicle were split in two groups of 3 mice each and injected with either buprenorphine (1 mg/kg, s.c.) or vehicle, and withdrawal thresholds assessed again 60 mins later. The following day after washout, the process was repeated with mice crossed over to the corresponding treatment.

Withdrawal thresholds were measured with the investigator blinded to treatment. Paw width was also measured from the ventral to dorsal aspects across the widest part of the paw using a digital micrometer before injection of CFA and then afterwards as an index of inflammatory oedema/load.

Cisplatin-induced neuropathic pain: CR (CD-1) female mice were purchased from Envigo and maintained at the animal facility at the University of Texas at Dallas. Mice were group-housed (4 maximum) in cages with bedding material provided for enrichment with food and water available *ad libitum* in a 12:12 h light-dark cycle. Room temperature was maintained at 21-22°C. Behavioral experiments were performed on mice 8-12 weeks old at the start of the experiment. Cisplatin is an alkylating agent that interferes with DNA crosslinking inside cells to induce cell cycle arrest and death. Due to the lack of a blood brain barrier, cisplatin (and other chemotherapeutics such as taxanes, vinca alkaloids) can accumulate in the DRG wherein it can orchestrate pathological changes to induce neuropathy as a dose-limiting side effect. Accordingly, we adopted a well characterized protocol for inducing signs of neuropathic pain in mice using repeated injections of cisplatin administered via 2 separate cycles^89^.

Once the presence of mechanical hypersensitivity had been established in cisplatin administered mice by Day 16, the mice were distributed into 2 separate treatment groups for administration of recombinant mouse meteorin (AF3475, R&D systems) at 1.8 mg/kg (s.c.) or vehicle at Days 18, 20, 22, 24, and 26. Withdrawal thresholds were measured pre and post cisplatin administration, and pre and post meteorin administration at the days indicated in Figure 2. On Day 34, three mice per group were euthanized to harvest tissue for immunohistochemistry with withdrawal thresholds routinely assessed in the remaining mice until they had resolved back to a baseline level.

Von Frey testing of mechanical allodynia: For SNI and CFA experiments, the mice were placed on a wire mesh in test containers for habituation (15-30 min habituation prior to testing). von Frey filaments (Stoelting, 0.02–2.0 g) were applied in ascending order to the lateral part (SNI) or mid plantar surface (CFA) of the hind paws. Each von Frey filament was applied five times and a positive response in at least three out of five stimuli determined the threshold level. A positive response was defined as sudden paw withdrawal, flinching, and/or paw licking induced by the filament^21,88,90^. Von Frey testing in cisplatin mice was initiated after habituating mice to acrylic behavioral chambers for 2 hrs. Testing was performed using the up-down method ^91^ with von Frey filaments applied to the mid plantar surface of the left hind paw. A positive response was comprised of licking or immediate flicking of the hind paw upon application of the filament.

### Immunohistochemistry (IHC)

Cisplatin administered mice were anesthetized using 4% isoflurane and euthanized by decapitation. DRG and skin tissues were flash-frozen in Optimum Cutting Temperature medium (Fisher Scientific) on dry ice. Twenty micron sections were cut on a cryostat and mounted onto SuperFrost Plus slides (Thermo Fisher Scientific). The sections were fixed in ice-cold 10% formalin for 15 min followed by incubation in an increasing percentage of ethanol 50%, 70%,100% for 5 min each. The fixed slides were transferred into blocking solution (10% Normal Goat Serum, 0.3% Triton-X 100 in 0.1 M phosphate buffer (PB)) for 1 hour at room temperature. Sections were incubated in primary antibody dissolved in blocking solution for 3 hours at room temperature or 4°C overnight. They were then washed in 0.1 M PB followed by incubation in secondary antibody diluted in blocking solution for 1 hour at room temperature. The slides were washed with 0.1 M PB followed by incubation with DAPI diluted in blocking solution for 5 min at room temperature. Lastly, the sections were washed in 0.1 M PB and cover-slipped using Prolong Gold Antifade (Thermo Fisher Scientific). Images were taken using an Olympus FluoView 3000 confocal microscope and analyzed using Cellsens (Olympus) software. All IHC images are representative of a sample size of 3 animals per group.

Image analyses of the DRG sections were performed by drawing a region of interest around individual neurons and the mean grey intensity (MGI) value was calculated in the targeted channel of interest. Average MGI was subtracted from the background fluorescence intensity calculated using the negative control (secondary antibody only without primary antibody) and normalized over the area surrounding each individual neuronal cell body.

### RNAscope and Integrated Co-detection Workflow (ICW)

ICW is a method that combines immunofluorescence with *in situ* hybridization, enabling simultaneous detection of proteins and nucleic acids. The ICW assays were performed in accordance with guidelines provided in the user manual for RNAscopeTM Multiplex Fluorescent Reagent v2 Assay combined with Immunofluorescence - Integrated Co-Detection Workflow (ACD, MK 51-150) for fixed-frozen tissue with the following specifications and adjustments: The tissue (DRGs from 12 weeks old, wild-type C57BL/6j mice) was fixed and post-primary fixed in 4 % PFA in 1X PBS. Antibodies were diluted in 4 % donkey serum in 1X PBS-T rather than Co-detection Antibody Diluent. Target retrieval was performed by submerging the slides in boiling 1X Co-Detection Target Retrieval Agent (ACD, 323165) solution for five min. Incubation with protease IV (ACD, 322331) for 30 min at 40°C in a humified chamber was chosen as the preferred protease treatment method. The sections were incubated with secondary antibodies for 2 hours. The following combination of probes, antibodies, HRP reagents, and fluorophores were used to examine meteorin expression in the DRGs: Goat anti-FABP7 (R&D Systems, AF3166-SP, dilution: 1:25) and donkey anti-Goat IgG, Alexa Flour 647 (Thermo Fisher, A21447, dilution: 1:300). Probes: Mm-Meteorin-C1 (mouse) (ACD, 131354, dilution: 1:1). HRP reagents: RNAscope Multiplex Fl v2 HRP-C1 (ACD, 323104). Fluorophores: TSA VividTM fluorophore 520 (ACD, 323271, dilution: 1:1500)

### DRG dissection and SGC culturing

Lumbar DRGs were isolated from post-natal day-3 mice into ice-cold HBSS (#14025092, Thermo Fisher) and centrifuged at 500xg for 4 min at room temperature. DRGs were then enzymatically dissociated by sequential 30-min incubations at 37 °C in papain (20 U/mL, Worthington #LS003126) and collagenase/dispase II (1 mg/mL each; Worthington #LS004176, Sigma #D4693); enzymes were removed by centrifugation (500xg, 1 min) and a wash in DMEM/F-12 (Sigma #D8437) supplemented with 10 % FBS and 1 % penicillin/streptomycin. The pellet was resuspended in the same medium containing DNase I (0.02 mg/mL), mechanically dissociated by gently trituration, filtered through a 100 µm cell strainer, and then the DRG cell suspension was pre-plated for 90 min at 37 °C/5 % CO₂ (2 wells of a 12-well plate) to enrich for adherent cells (enriched for SGCs). Media containing non-adherent cells was then carefully removed and fresh media added. Cells were maintained at 37 °C/5 % CO₂ with medium changes every 48 hour.

### SGC Stimulation assay

On Day 6 cultures were serum-deprived overnight to suppress basal phosphorylation. The next day cells were stimulated with recombinant mouse Meteorin (800 ng/mL; R&D Systems), or received vehicle only, for 15 at 37 °C/5 % CO₂. Stimulations were terminated simultaneously on ice by two rapid PBS rinses and lysis in ice-cold TNE-buffer (1% Nonidet P40, 10 mM Tris base, pH 8, 150 mM NaCl, 1 mM EDTA) containing Complete ULTRA mini (Roche, 05892791001) and Phos-STOP inhibitors (Roche, 4906845001). Cell lysates were rotated for 30 min at 4 °C, clarified (14 000 xg, 5 min, 4 °C) and total protein quantified by BCA assay. Equal amounts of protein (30 µg) from each condition was mixed with NuPAGE sample reducing agent (Invitrogen, NP0009) and NuPAGE LDS 4x Sample Buffer (Invitrogen, NP0007), resolved by SDS-PAGE on 4–12 % Bis-Tris gels (Invitrogen, NP0322BOX) in MOPS SDS Running Buffer (Invitrogen, NP0001), and transferred to nitrocellulose using an iBlot 2 dry-blot stack (Invitrogen, IB23002). Membranes were blocked for 1 h in BSA blocking buffer, probed overnight at 4 °C with rabbit phospho-specific antibodies to p44/42 MAPK (ERK1/2) (Thr202/Tyr204) (Cell Signaling Technology, 9101) and Akt (Thr308) (Cell Signaling Technology, 2965), or mouse anti-GAPDH (Sigma, G8795), washed, and incubated with HRP-conjugated secondary antibodies for 1.5 h at room temperature. Chemiluminescence was developed with ECL reagent (Cytiva, RPN2109) and imaged on an iBright 1500 platform; exposure times were kept within the linear range. Band intensities were quantified in iBright Analysis software with background subtraction, normalised to GAPDH, and expressed relative to vehicle-treated samples for statistical evaluation.

### Calcium imaging

Transcardial perfusion was performed using 20 ml DPBS (Thermo Scientific, SH3002802) on mice anesthetized with isoflurane (IsoFlo Vet, Abbott). All DRGs were collected and kept in ice-cold HBSS (14170088, Thermo Fisher). The tissue was centrifugated at 500xg for 4 min at room temperature. Sequentially the tissue was incubated with collagenase (1.25 mg/ml; C9722, Sigma Aldrich) and dispase II (2.5 U/ml; D4693-1G, Sigma Aldrich) dissolved in 2 ml DPBS (Thermo Scientific, SH3002802) for 30 min at 37°C, 5% CO2. The cells were resuspended in 1ml DMEM (v/v) with 10% Fetal Bovine Serum, 1% Penicillin and 1 μl DNase Cells were gently triturated and mechanically dissociated until the solution gained a homogenous aspect, and 9 ml DMEM was added. The cells were centrifugated at 500xg for 8 min at room temperature. The pellet was resuspended in 4-6 ml of DMEM, 10% FBS, 1% Penicillin and plated in wells coated overnight with laminin (L2020, Sigma Aldrich) and 2h poly-L-lysine (0.05mg/ml; 8920, Sigma Aldrich). After 18 hours of incubation, DRG primary culture was loaded with 1 μM Fura 2 AM (F1221, Thermo Fisher) in HBSS (HBSS with 10 mM HEPES without phenol red, Stem cell) for 45 min in the incubator and protected from light. Sequentially the samples were incubated for 15 min in calcium-enriched HBSS media. Fura 2 AM primarily stained SGCs allowing the visualization of these cells. Cells were washed for 15 min before the experiment initiation to avoid any Acetoxymethyl ester left to promote Fura 2 AM penetration in intracellular compartments. Calcium imaging was conducted at a closed bath imaging chamber (RC-21BRFS, Warner Instruments) with a constant buffer (HBSS with 10 mM HEPES without phenol red, Stem cell) flow of 800 μl/min (Infuse/withdraw pump II pico plus elite, Harvard Apparatus). The samples were attached to the chamber within the coverslip (CS-25R, Warner Instruments), where it was previously incubated, and the chamber was screwed in the microscope support to avoid any movement. The addition of stimulants occurred manually through the injection port at the imaging chamber while the buffer flow was paused. Samples were stimulated with recombinant mouse meteorin (200-800 ng/ml, AF3475, R&D systems), ATP (10 μM, Thermo Fisher), Ionomycin (1 μM; Alomone), or Endothelin 1 (1.6 nM; ET1, Sigma Aldrich) diluted into flow buffer as described above. All imaging analyses were done with an inverted wide-field fluorescence microscope (10x lens, Olympus IX81) at 37°C and a 5% CO2 atmosphere. Cell sens-vision software was used for data analysis, and Camera Prime 95B (Quantum Efficiency of 95%, Tiledyne Photometrics) was used for recording. Samples were excited at 340 nm and 380 nm wavelength, presenting emission at 510 nm wavelength. Images were recorded every 300 ms. The fluorescence ratio of 340 nm/380 nm indicated a relative change in the cytoplasmatic calcium concentration in the selected region of interest.

Data analysis: A positive calcium signal was defined as a response 20% above baseline after exposure to a stimulant. After background subtraction, it was determined the average of the last five seconds of readings from the bath treatment before a stimulant addition. The maximum value registered after the stimulant treatment was also determined. Finally, the percentage signal is determined, % [Ca2+]i =Maximum-Average/Average.

### Mass spectrometry

DRGs were isolated from 12 wild type C57Bl/6JRj mice, 6 female and 6 males. Cell cultures were prepared as described above and seeded in the following manner for paired stimulation experiment: The DRGs from 2 mice (1 female and 1 male) were collected in one tube, dissociated as described, and seeded in 2 wells in a 12-well plate (1 mouse/well). One of the respective wells was stimulated with 200 ng/mL recombinant mouse meteorin (3475-MN, R&D Systems) at time zero and another dose after 24 hours. The control well was incubated with corresponding volumes of PBS.

Samples for proteomics analyses were prepared 48 hours after first stimulation by washing the cells in preheated PBS and lysing in 200 ul lysis buffer (5% sodium dodecyl sulfate (SDS), 5 mM tris(2-carboxyethyl)phosphine (TCEP), 10 mM chloroacetamide (CAA), 100 mM Tris, pH 8.5). Lysates were transferred to Eppendorf tubes, boiled for 10 min at 95°C, and sonicated with a micro tip probe. Protein concentrations were estimated using the Pierce BCA Protein Assay Kit (Thermo Fisher Scientific). For each sample all protein (approx. 300-450 µg) was used for PAC digestion in an automated 96-well format on a KingFisher™ Flex robot (Thermo Fisher Scientific) with 12-hour overnight digestion at 37 °C using LysC (Wako) and trypsin (Sigma-Aldrich) as described in Bekker-Jensen et al., 2020^92^. Protease activity was quenched by acidification with trifluoroacetic acid (TFA). Peptides were purified and concentrated on reversed-phase C18 Sep-Pak cartridges (Waters) and eluted with 40% acetonitrile (ACN) followed by 60% ACN. The eluate was concentrated, and ACN removed using a SpeedVac centrifuge, and the peptide concentration was estimated by measuring absorbance at A280 on a NanoDrop (Thermo Fisher Scientific). Identical amounts of each sample were labelled using TMT11-plex reagents (Thermo Fisher Scientific) according to the instructions of the manufacturer. Samples were mixed and purified on Sep-Pak, concentrated in a SpeedVac centrifuge, and fractionated by microflow high pH reversed-phase fractionation on an Ultimate 3000 HPLC system (Dionex) using a Waters Aquity CSH C18 1.7 μM 1 x 150 mm column operating at a constant flow rate of 30 μl/min. Buffer A (5 mM ammonium bicarbonate (pH 8)) and buffer B (100% acetonitrile) were used to separate the peptides into 46 fractions collected without concatenation. All fractions were acidified by addition of 10% formic acid, dried in a SpeedVac centrifuge and resolubilized with 0.1% formic acid before loading on Evotips prior to MS analysis. Samples were analyzed on the EvoSep One system (using the pre-programmed 60 samples per day gradient) coupled to an Orbitrap Exploris 480 MS (Thermo Fisher Scientific) through a nanoelectrospray source. Peptides were separated on a 15-cm, 150 μM inner diameter analytical column in-house packed with 1.9 μM reversed-phase C18 beads (ReProsil-Pur AQ, Dr Maisch) and column temperature was maintained at 60°C by an integrated column oven (PRSO-V1, Sonation GmbH). Proteome analysis was performed using data-dependent acquisition (DDA). All raw files were analyzed using MaxQuant software version 1.6.0.17 with the integrated Andromeda search engine^93,94^. Files were searched against the mouse UniProt database, supplemented with commonly observed contaminants. Data analysis of proteomics data was performed using Perseus software version 1.6.5.0. Data were log2 transformed and filtered. A minimum of three valid values were required for a protein identification to be included in downstream analysis. Data were normalized by quantile-based normalization. One of the PBS control samples were considered an outlier based on initial quality control assessment of the data and excluded from the downstream analysis. Data was normalized by row-based median subtraction within groups and significantly regulated proteins comparing meteorin and vehicle treatment were identified by t-test using a significance cut-off of 0.05 and S0=0.1. Heatmaps were generated based on unsupervised hierarchical clustering. Volcano plots were generated for visualization of significantly regulated proteins identified by Student’s t-test (significance cut-off FDR<0.05 and S0=0.1). The plots show the –log10-transformed p-values against the log2 fold changes comparing meteorin and PBS treatment, and significantly regulated proteins were determined based on a hyperbolic curve threshold generated using the described statistical parameters. **Supplementary table 1** includes all identified proteins with statistically significant regulation indicated by a “+” in the first column (Table S1). Gene ontology (GO) term enrichment analysis and Kyoto Encyclopedia of Genes and Genomes (KEGG) pathway enrichment analysis were done using InnateDb^95^.

### Intracellular Luficer yellow Labeling

Experiments were performed on Balb/c mice of either sex (M:F 1:1), aged 2-5 months. At Day 18, control and SNI mice were sacrificed by CO_2_ inhalation, and DRGs L4,5 were removed and placed in cold (4°C) Krebs solution (pH 7.4) containing (mM): 120.9 NaCl, 4.7 KCl, 14.4 NaHCO3, 2.5 MgSO_4_, 1.2 NaH_2_PO_4_, 2.5 CaCl_2_, and 11.5 glucose. The ganglia were pinned onto the silicon rubber-covered bottom of a chamber superfused with Krebs solution bubbled with 95% O_2_ and 5% CO_2_ at 23–24°C.

The chamber was placed on the stage of an upright microscope (Axioskop FS, Zeiss, Jena, Germany) equipped with fluorescent illumination, x40 water immersion lens, and a digital camera (Penguin 600CL, Pixera Corp., Los Gatos, CA, USA) connected to a PC. Single SGCs were injected with the fluorescent dye Lucifer yellow (LY, Sigma-Aldrich, 3% in 0.5 M LiCl) from a glass microelectrode connected to a preamplifier (Neuro Data Instrument Corp., New York, NY, USA). Tip resistances of microelectrodes were 80-120 MΩ, and the dye was injected by hyperpolarizing current pulses, 100 ms in duration and 0.5 nA in amplitude at 5 Hz for 3–5 min. During and after injections, the LY-labeled cells were imaged with the camera. The number of SGCs coupled to the LY-injected cell was counted shortly after the dye injection (see ^44^). SGCs were considered as coupled when the dye passed from SGCs around a given neuron to SGCs around one or more neighboring neurons. Results from SNI and control mice were compared. Dye coupling data were pooled for each time point from multiple experiments; This was done because relatively small numbers of cells were injected per experiment. When an LY-injected cell was found to be dye coupled it was marked as 100, and when it was not coupled, it was marked as 0, as previously described^96^. These data were analyzed using one-way ANOVA with Tukey’s multiple comparison test.

### Data and Statistical Analysis

SNI, cisplatin and CFA mechanical thresholds: Data analysis was performed using Graphpad Prism 8.4.1. Statistical differences between groups were assessed using two-way repeated measures (RM) ANOVA when every subject had data at every time point. When values were missing a mixed model ANOVA was applied. In each case this was followed by Tukey’s test. AUC effect sizes were determined by subtracting behavior scores from corresponding baseline values for each animal at all timepoints displaying significant differences with the designated control group. These values were summed and expressed as a % of the designated control group and compared by one-way ANOVA followed by Tukey’s test. All data are represented as mean +/- SEM with p<0.05 considered significant.

Calcium influx assay: Correlation between groups were analyzed using Chi-square and Fisher’s exact test and only p value < 0.05 were considered significant. For Chi-square analysis, significance was reached with p-value, p < 0.0001, p < 0.0126. For Fisher exact test, significance was reached with p value, p < 0.0001, p =0.0005, p =0.0078.

---

**Supplementary Table 1.**

MS raw data file

**Supplementary Figure S1.**
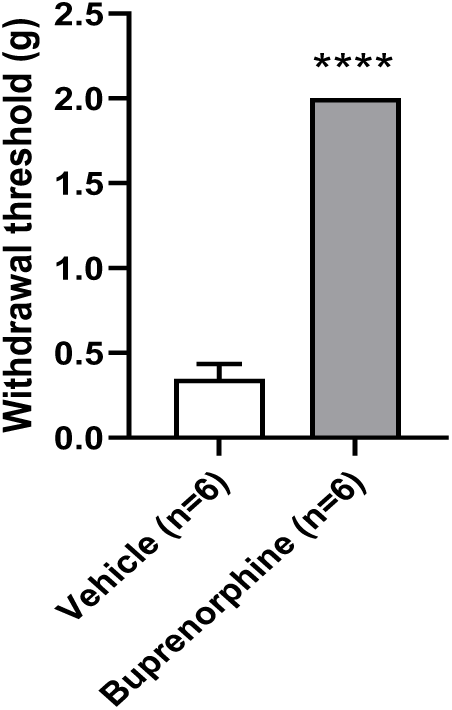
Buprenorphine treatment. Upon completion of the experiment in CFA mice assessing meteorin efficacy (meteorin vs vehicle), 6 CFA administered female mice treated with repeated injections of vehicle were treated with either the partial m-opioid receptor agonist buprenorphine (Temgesic®; 0.1 mg/kg, s.c., n=3) or Vehicle (PBS s.c., n=3). Effects on mechanical withdrawal thresholds (g) to von Frey stimulation were then assessed. The following day, the treatments within the two groups were switched and effects on mechanical withdrawal thresholds (g) to von Frey stimulation assessed once again. This crossover paradigm enabled 6 mice in each treatment group to be obtained. For the mechanical assessment, 2 grams were cut-off, and this value was reached for all mice in the buprenorphine group, The experimenter was blinded to treatment. ****P<0.0001 vs. Vehicle, Student’s t test. Data are means ± SEM.

**Supplementary Figure S2.**
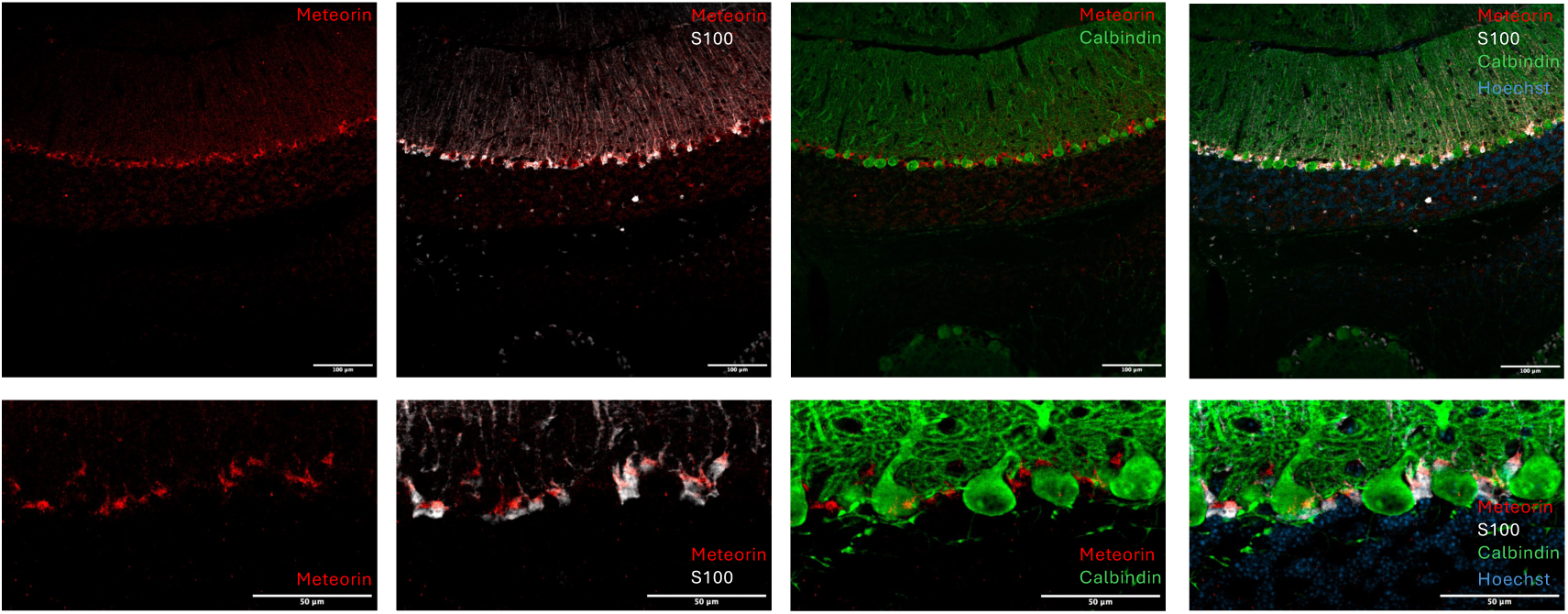
Meteorin expression in mouse cerebellum Bergmann glia. Visualization of meteorin by IHC of mouse cerebellum, showing staining af Bergmann glia. Confocal microscopy of cerebellum stained with antibody against meteorin (red. R&D Systems, AF3475), the glial marker S100 (white. Dako, GA504), the Purkinje cell marker calbindin (green. Novus Biologicals, NBP2-50028), and Hoechst (blue). Top: 10x magnification. Bottom: 40x magnification.

**Supplementary Figure S3.**
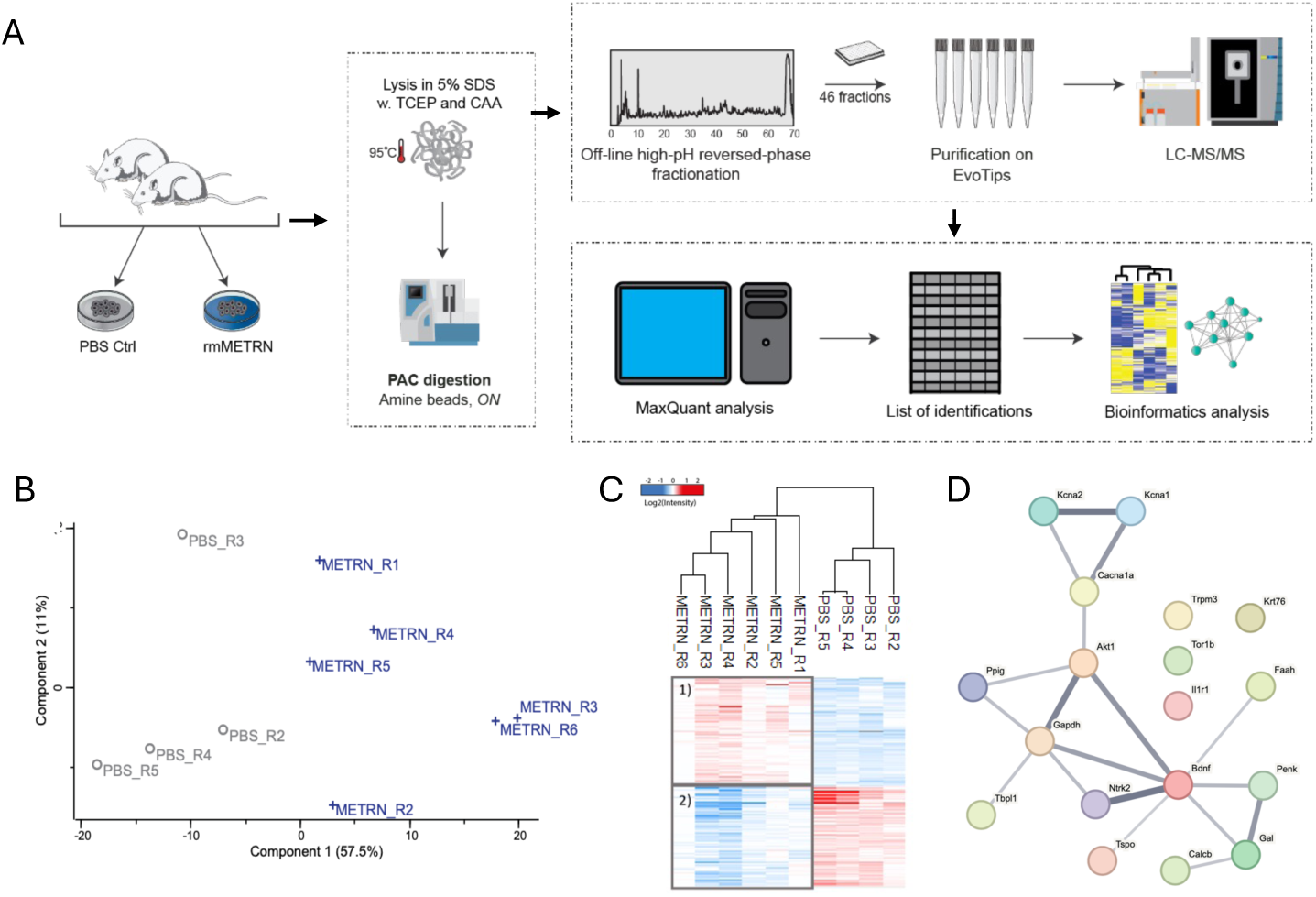
Global proteome changes of meteorin stimulated DRG cultures. **A)** Experimental design: Proteome analysis of mouse DRG cultures incubated with or without meteorin for 48 hours. Schematic workflow depicting the different steps underlying the MS-based proteome analysis. These include the experimental set-up incorporating “paired” PBS controls and meteorin stimulation, sample lysis and digestion, off-line fractionation followed by loading on Evotips and LC-MS/MS analysis, and finally raw data processing and bioinformatics analysis. **B)** PCA plot shows that samples cluster according to treatment (PBS_R2-6, METRN_R1-6). **C)** Identified proteins cluster according to treatment. Heatmap based on unsupervised hierarchical clustering of all identified proteins. Color scale indicates normalized log2 intensities. Samples: PBS_R2-6, METRN_R1-6. **D)** Unbiased identification of regulated proteins implicated in pain. Analysis resulted in a list of 18 proteins displayed in a STRING-based network.

**Supplementary Figure S4.**
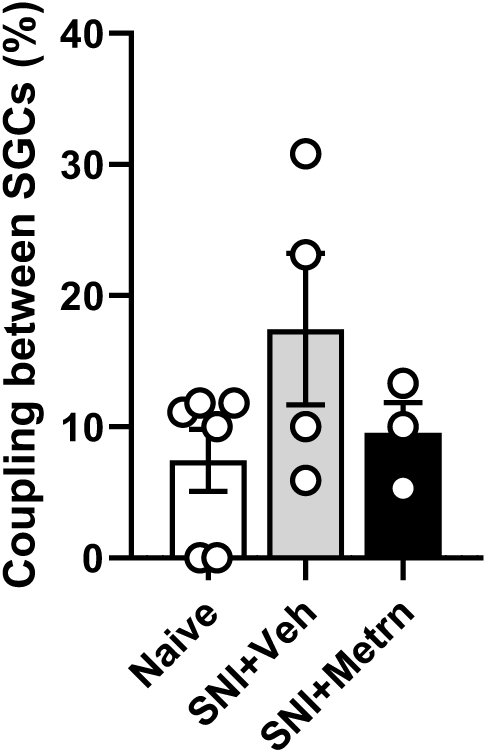
Meteorin reverses signs of injury-induced pathophysiology at DRG cell bodies. SNI increases SGC coupling and is reversed by meteorin treatment. Data are shown as means ± S.E.M.

